# Structural landscape of the complete genomes of Dengue serotypes and other viral hemorrhagic fevers

**DOI:** 10.1101/2020.04.23.056986

**Authors:** Riccardo Delli-Ponti, Marek Mutwil

## Abstract

**Background:** With more than 300 million potentially infected people every year, and with the expanded habitat of mosquitoes due to climate change, dengue cannot be considered anymore only a tropical disease. The RNA secondary structure is a functional characteristic of RNA viruses, and together with the accumulated high-throughput sequencing data could provide general insights towards understanding virus biology. Here, we profiled the RNA secondary structure of >7500 complete viral genomes from 11 different species of viral hemorrhagic fevers, including dengue serotypes, ebola, and yellow fever.

**Results:** We achieved hig prediction scores (AUC up to 0.85 with experimental data), and computed consensus secondary structure profiles using hundreds of structural *in silico* models. We observed that virulent viruses such as DENV-2 and ebola tend to be less structured than the other viruses. Furthermore, we observed virus-specific correlations between secondary structure and the number of interaction sites with human proteins, reaching a correlation of 0.89 in the case of zika. We demonstrate that the secondary structure and presence of protein-binding domains in the genomes can be used as intrinsic signature to further classify the viruses. We also used structural data to study the geographical distribution of dengue, finding a significant difference between DENV-3 from Asia and South-America, which could imply different evolutionary routes of this subtype.

**Conclusions:** Our massive computational analysis provided novel results regarding the secondary structure and the interaction with human proteins, not only for Dengue serotypes, but also for other viral hemorrhagic fevers. We also provided a new approach to classify viruses according ot their structure, which could be useful for future cassifications. We envision that these approaches can be used by the scientific community to further classify and characterise these complex viruses.

## BACKGROUND

Dengue is a mosquito-borne virus that can potentially infect more than 300 million people a year in more than 120 countries ^1,2^. Dengue infection can further evolve into a severe hemorrhagic fever, which could lead to shock and death. Due to climate change, the disease is now threatening an increasing number of countries, with cases reported in Europe ^2^. The existence of four different serotypes (DENV-1, 2, 3, 4), with also a fifth recently reported ^3^, complicates the development of an effective vaccine ^4^. The four serotypes show not only significant differences in sequence similarity ^5^ but also distinctive infection dynamics. For example, DENV-1 is the most wide-spread serotype, followed by DENV-2 ^6,7^, which is also more often associated with severe cases ^8^. However, the mechanisms behind Dengue infections and the complete set of differences between the serotypes are still unclear.

Dengue is only one of the different Viral Haemorrhagic Fevers (VHF), comprising single-stranded RNA viruses from different families, such as flavivirus and filovirus. These viruses can be extremely lethal in humans, as in the case of Ebola. Other VHFs show not only similarities to Dengue in terms of symptoms, but also in the transmission vector. For example, Yellow Fever is also a mosquito-borne VHFs, with higher mortality but a slower rate of evolutionary change compared to Dengue ^9^. The mild VHF Chikungunya shares the same vector with Dengue, mosquito *Aedes aegypti*, and the two viruses can even coexist in the same mosquito ^10^. However, even with thousands of viral genomes available, the most common approach to understand the similarities between those viruses is still based only on the sequence.

The secondary structure of RNA viruses is fundamental for many viral functions, from encapsidation to egression from the cell and host defence ^11–13^. Specific structures in the UTRs were found to be functional, for example, in Dengue, but also in HIV and coronaviruses ^11,14,15^. Other structural regions, including the 3’ UTR, were found conserved not only in Dengue serotypes but also between Dengue and Zika ^16^. Moreover, the single-stranded RNA (ssRNA) viruses preserve their structure (folding) even if their sequence mutates rapidly ^17,18^. Thus, the folding shows the potential to be used to classify different viral species and subspecies.

However, while the RNA secondary structure is an informative element to characterise viruses, the secondary structure of only a few viral genomes has been experimentally characterized ^16,19^. Consequently, while thousands of viral genomes have been sequenced, we can only rely on *in silico* data to study their secondary structure. Furthermore, predicting the RNA secondary structure of entire viral genomes can be challenging, due to usually large sizes of >10000 nucleotides (nt), where most thermodynamic algorithms used to model the secondary structure drop in performance after 700 nt ^20^. In our work, we computationally profiled the RNA secondary structure of >7500 viral genomes (prioritising Dengue serotypes and in general VHFs) using Computational Recognition of Secondary Structure (CROSS), a neural network trained on experimental data, which was successfully applied to predict HIV genome structure ^21^. We mapped the secondary structure properties of the viruses on the world map, to study the genome interaction with proteins, and to further classify and understand the viruses.

## METHODS

### Source of viral genomes

The viral genomes were downloaded from NCBI, selecting for each specific virus the fasta sequences containing the keyword *complete genome*. NCBI data was also used to extract the geographical information of Dengue viruses as well as the serotypes. Fragmented or incomplete genomes passing the filter were also removed by looking at the sequence length.

### RNA Secondary structure

The secondary structure profiles were computed using the CROSS (Computational Recognition of Secondary Structure) algorithm. CROSS is a neural network-based machine learning approach trained on experimental data, able to quickly profile large and complex molecules such as viral genomes without length restrictions. CROSS was already used to profile the complete HIV genome, also showing an AUC of 0.75 with experimental ‘Selective 2′ Hydroxyl Acylation analyzed by Primer Extension’ (SHAPE) data ^21^. For a comprehensive analysis, we used the *Global Score* model, considering nucleotides with a score >0 as double-stranded, and <0 as single-stranded. For the purpose of computing the structural content of a complete genome (i.e., % double-stranded nucleotides), the total number of nucleotides with a score >0 was averaged for the total length of each genome. For the plots showing the complete secondary structure of dengue regions, we used the MFE structure computed using RNAfold ^22^.

### Protein-RNA interactions

To analyse the protein binding motifs in the viral genomes, we selected 5-mer motifs from the table S3 of Dominguez et al. ^23^. All of the 520 possible redundant motifs (270 non-redundant) were selected for further analysis. We scanned for the motifs on the complete genomes of the different strains, selecting only perfect matches. The number of motifs normalized by the average genome length of the different species of viruses was used to define a score for the number of potential interactions with proteins, according to the following formula:

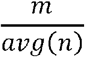

where m is the number of exact motifs found in a genome, and avg(n) is the average length of the genome of a specific species.

We also computed high-throughput predictions against the human proteome using the catRAPID Omics algorithm ^24^, which estimates the binding propensity between proteins and RNA by combining secondary structure, hydrogen bonding and van der Waals contributions. The algorithms computed more than 2 millions interactions between viral genomes and human proteins. The Discriminative Power (DP) was used to progressively filtering for strong interactions. The Discriminative Power (DP) ranges from 0 to 1, where DP values above 0.5 indicate that the interaction is likely to take place.

### Hierarchical clustering

The structural content and the averaged number of binding domains were employed to build dendrograms. To this end, we computed the Euclidean distance between the values associated with each virus using statistical software R. We then used the *hclust* function based on a *ward*.*D2* module to build the dendrograms according to the hierarchical clustering.

### Secondary structure consensus profile

We used hundreds of profiles generated using CROSS to build secondary structure consensus profiles for Zika and Chikungunya. To build the consensus profiles we selected non-overlapping windows of 50 nucleotides across all the genomes of each species, and we averaged CROSS propensity scores for each window in all the genomes available. To avoid problems due to the different lengths of the genomes, we limited the sliding window till the average length of the genomes of that specific species (Table 1). Regions with a score >0 are double-stranded regions in agreement in all the genomes of a species, while <0 for single-stranded consensus regions.

**Table 1.** The information regarding the number of genomes available, the family, and the average nucleotide length of each family for all the viruses used in our analysis.

| Genome | Family | Hemorrhagic | Number of genomes | Avg Size |
| --- | --- | --- | --- | --- |
| Denv-1 | Flavivirus | Yes | 1653 | 10500 |
| Denv-2 | Flavivirus | Yes | 1322 | 9750 |
| Denv-3 | Flavivirus | Yes | 775 | 10590 |
| Denv-4 | Flavivirus | Yes | 192 | 9730 |
| Zika | Flavivirus | No | 274 | 10120 |
| Chikungunya | Togavirus | Yes*<br>(mild) | 585 | 10500 |
| Japanese encephalitis | Flavivirus | Yes*<br>(rare cases) | 288 | 10500 |
| Yellow fever | Flavivirus | Yes | 158 | 8530 |
| West Nile fever | Flavivirus | Yes | 1660 | 10390 |
| Tick encephalitis | Flavivirus | Yes | 135 | 9770 |
| Ebola | Filovirus | Yes | 549 | 18200 |

## RESULTS

### Structural properties of the Dengue genomes

Here, we analysed, for the first time to our knowledge, the secondary structure profiles of the complete genomes of more than 7500 ss-RNA viruses (Table 1; Methods: Source of viral genomes). The structural profiles were generated using the CROSS algorithm, a fast and comprehensive alternative to profile the structural content (i.e., % of double-stranded nucleotides) of long and complex RNA molecules, such as viruses (^21^; see Methods: RNA secondary structure).

To analyse Dengue secondary structure, we selected one strain for each serotype, focusing on strains that were widely used in previous publications ^25^. In general, the four serotypes show significant differences in sequence, with around 65-70% sequence similarity ^5^. Their secondary structure also shows notable differences (Figure 1). For example, the 3’ UTR of DENV-1 shows a peculiar structural valley, compared to the others. Interestingly, DENV-1 and DENV-2 share the highest structural peak around 6000 nucleotides, while DENV-3 and DENV-4 also have the highest structural peak in common, but at position 4000.

**Figure 1.**
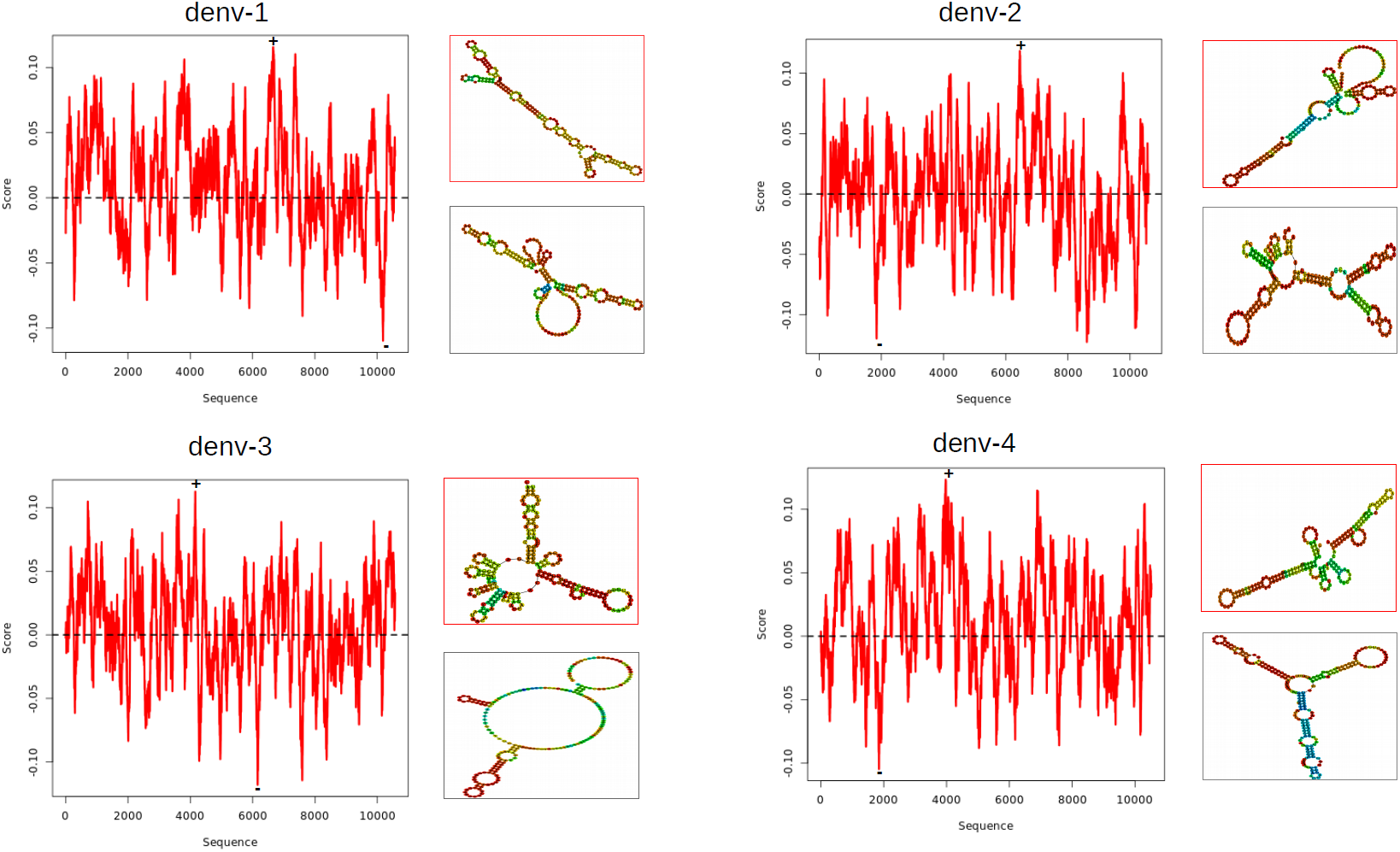
Secondary structure of the four dengue serotypes represented as propensity profiles. Nucleotides with a score >0 are double-stranded, while <0 indicates single-stranded nucleotides. The profile is normalised using the same formula reported in the original paper of CROSS methodology. For each profile, the highest (+) and lowest (-) structural peak is highlighted. The structures of 200 nt regions, including the most high-propensity double- (red) and single-stranded (gray) regions for each serotype, were computed using *RNAfold*.

We further expanded our analysis to cover the 4 different serotypes of the dengue virus, comprising ∼4000 genomes (Figure 2; Table 1). The analysis revealed that DENV-2 and DENV-3 are less structured than DENV-1 and DENV-4. It is well known that a lower amount of secondary structure can improve the translation efficiency ^26^, suggesting that DENV-2 and DENV-3 could be more efficiently translated in the host cell.

**Figure 2.**
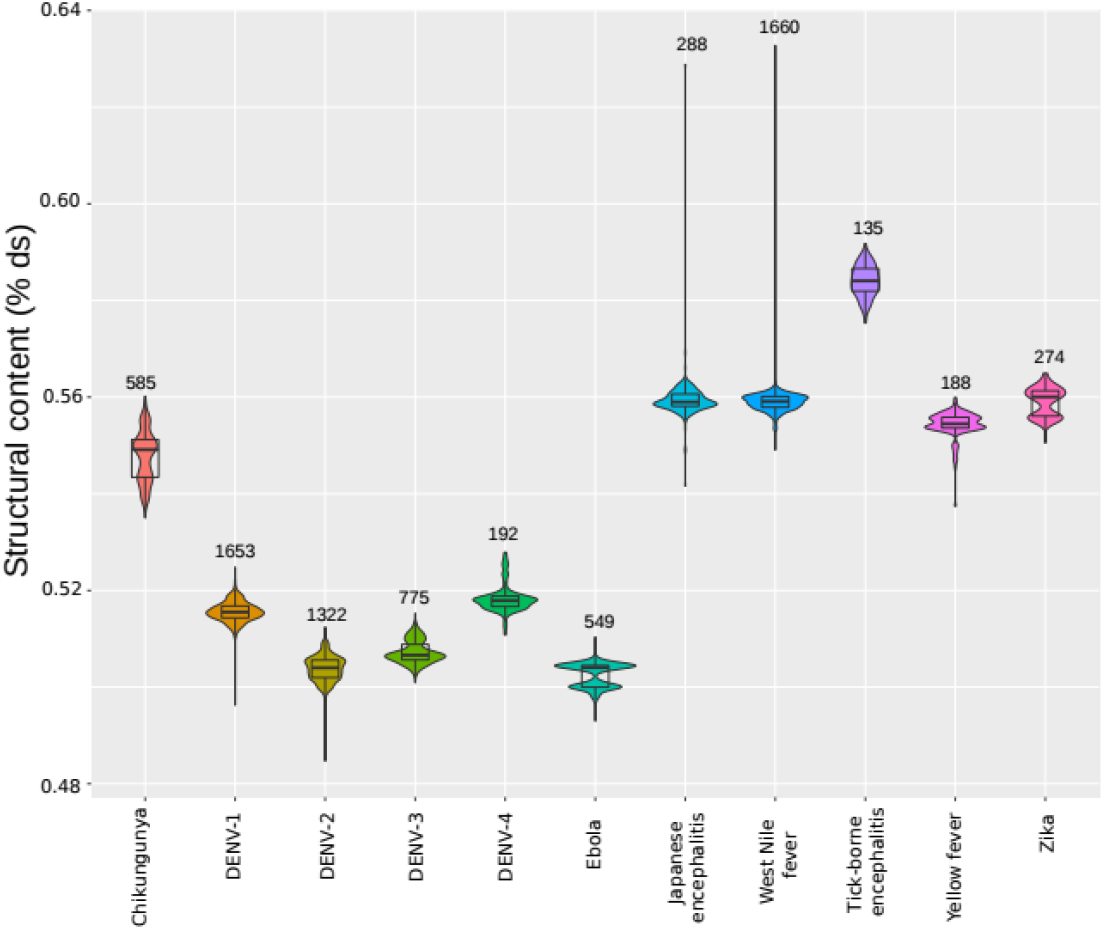
Structural content (% double-stranded nucleotides) for all the genomes for the 11 species. The number above each violin plot indicates the number of genomes used in each species.

To validate our approach, we compared our predictions with SHAPE experiments performed on Dengue genomes ^16^. Using the Area Under the ROC curve (AUC) to distinguish ranked SHAPE reactivities, we obtained an AUC ranging from 0.75 to 0.85 for DENV-2 and DENV-1 (Supplementary Figure 1a, b). This further proves the power of our in silico approach, which can generate thousands of secondary structure profiles with high performances on experimental data.

### Comparison of structural properties of the VHF genomes

Interestingly, dengue serotypes tend to be less structured than other flaviviruses, such as West Nile Fever, Yellow Fever, Tick-borne Encephalitis, and Japanese Encephalitis (Figure 2). Even if not properly a VHF, we also used as comparison Zika, due to the similarities with Dengue not only in the vector (*Aedes aegypti*), but also in terms of secondary structure domains ^16^. Interestingly, while Tick-borne Encephalitis and Zika genomes are more structured (average double-stranded nucleotides > 56%), Nile Fever and Japanese Encephalitis have a similar structural distribution, especially since they are also close in the species tree ^27^. To further compare and classify the secondary structure of other VHFs outside of flavivirus, we also included >500 genomes of Ebola and Chikungunya (Figure 2, Table 1). The analysis revealed that the other viruses are significantly more structured than Dengue (mean structural content for Flaviviruses and Dengue serotypes is 0.55, 0.51, respectively; Kolmogorov-Smirnov < 2.2e-16), with the exception of Ebola, which is not only the most deadly virus in our dataset (50% mortality rate according to WHO; ^28^), but also one of the less structured (mean structural content 0.50).

### Structural properties of the untranslated regions of VHF genomes

To further study the secondary structure content for the >7500 viral genomes, we also analysed the 5’ and 3’ UTRs (first 1000 nt considered 5’ UTR; last 1000 nt considered 3’ UTR; Figure 3a, b). DENV-3 is the only serotype with both UTRs more structured than the entire genome (5’ UTR= 0.55 and 3’ UTR=0.53; Figure 2), while DENV-1 has a more structured 5’ UTR (structural content = 0.53). This result is in line with the experimental Parallel Analysis of RNA Structure (PARS) data coming from human RNAs, where the UTRs were more structured than the CDS ^29^. This suggests that some viruses tend to mimic the secondary structure of human mRNAs to be efficiently translated by the cellular machinery ^30^. This is also further supported in Dengue, where a complex structure at the 3’ UTR was shown to mimic the absent polyA, to enhance translation ^31^. Interestingly, Ebola has the least structured UTRs (structural content 5’ UTR = 0.46; 3’ UTR = 0.41). In Zika, the 3’ UTR is more structured than the 5’ (Figure 3c; 3’ UTR=0.56, 5’ UTR = 0.50). Chikungunya shows not only the highest structural variability in the 3’ UTR (standard deviation 3’ UTR= 0.16, Figure 3b), with a more structured 5’ UTR (3’ UTR=0.43, 5’ UTR = 0.51; Figure 3a). Finally, Ebola, DENV-1, and DENV-3 exhibit a more structured 5’ UTR, especially when compared with DENV-2, DENV-4 and Japanese Encephalitis, which tend to be more structured (Figure 3c).

**Figure 3.**
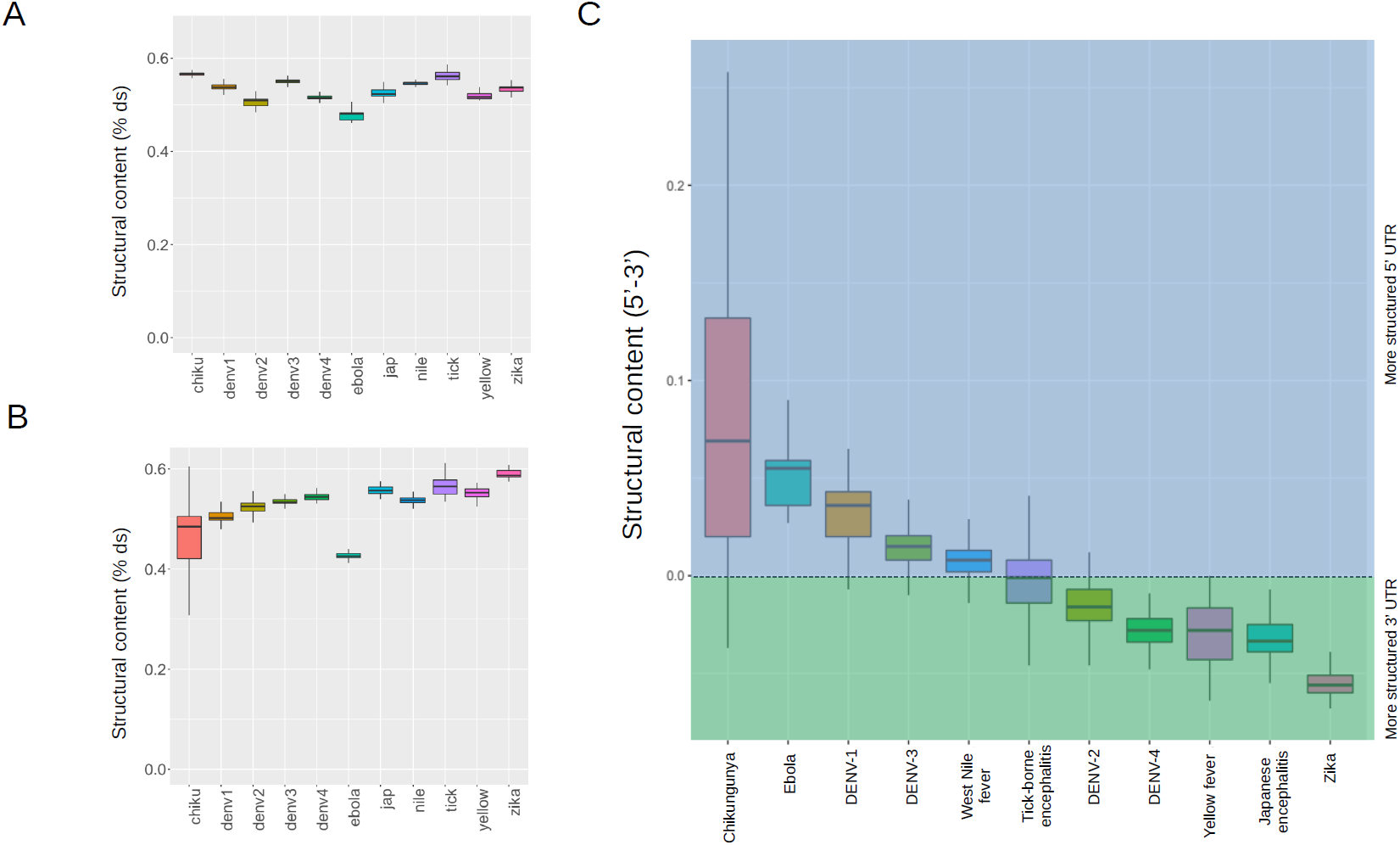
Structural content of the UTRs of the 11 viral species. (A) Structural content (% double-stranded nucleotides) for all the genomes for the 11 species for the 5’ UTR. To have an equal comparison between the different species, we considered the 5’ UTR as the first 1000nt. The abbreviations used for the viruses: Jap (Japanese encephalitis), Nile (West Nile Fever), Chiku (Chikungunya), Yellow (Yellow Fever) and Tick (Tick-borne encephalitis). (B) Structural content (% double-stranded nucleotides) for all the genomes for the 11 species for the 3’ UTR. To have an equal comparison between the different species, we considered the 3’ UTR as the last 1000 nt. (C) The difference for each individual genome between the structural content of the 5’ and the 3’ UTR. Viruses with more structured 5’ UTR are >0 (blue area), while <0 indicates more structured 3’ UTRs (green area).

### Structural content can be used to classify VHFs

The overall similarities and differences in structure are an additional feature that could be employed to characterise the different viruses. For the next step, the structural content (mean of the % of double-stranded nucleotides for all the viral genomes) in a specific species was used to hierarchically group the 11 different viruses (Methods: Hierarchical clustering; Table 1). The resulting dendrogram clustered the Dengue serotypes, showing that they are more structurally similar compared to other viruses (Figure 4). The structural similarities of Dengue serotypes, together with the similarity between West Nile Fever and Japanese Encephalitis, are in agreement with the Phylogenetic Tree of Viral Hemorrhagic Fever ^27^.

**Figure 4.**
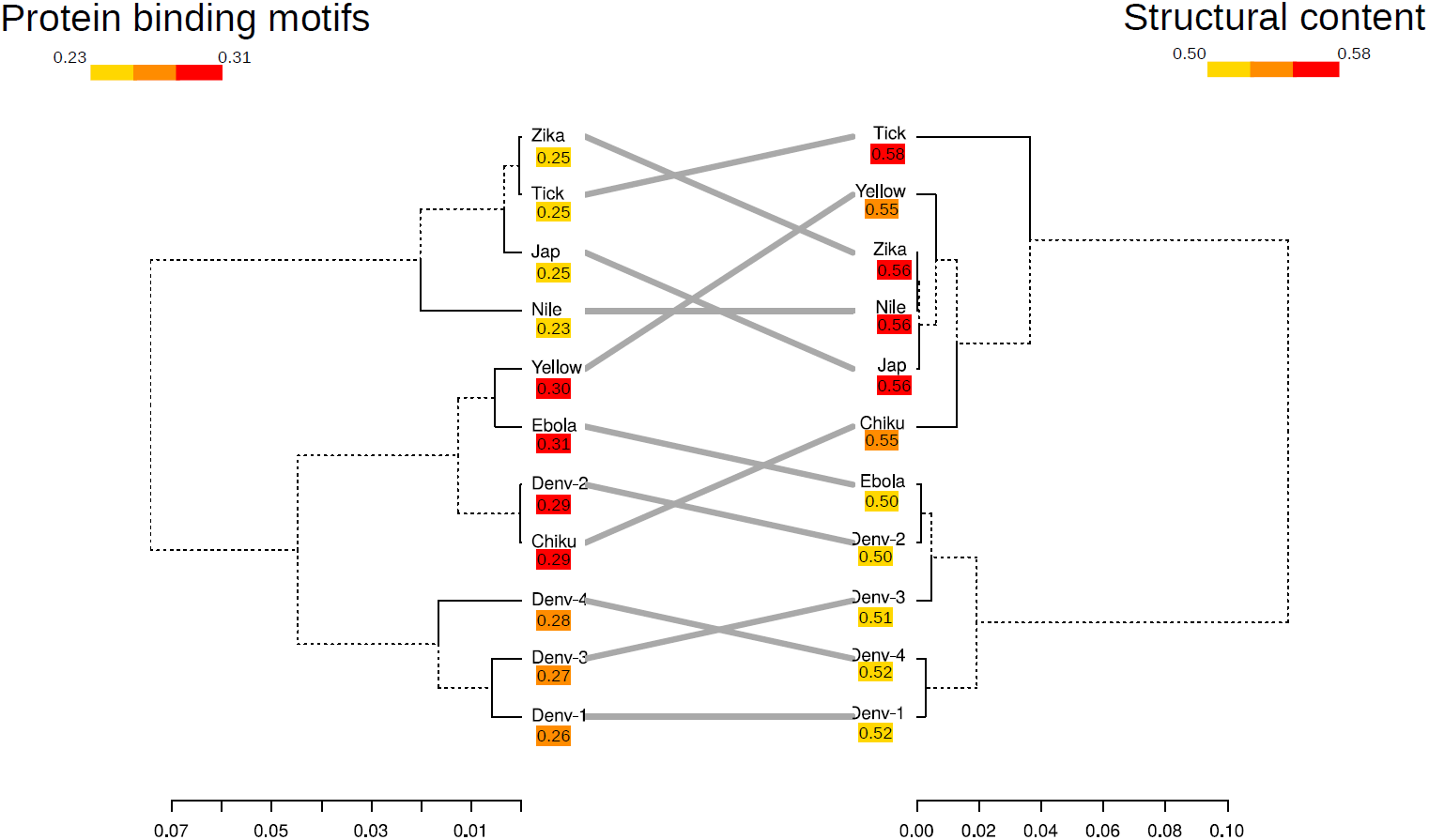
Comparison between the dendrograms obtained using the structural content (left side figure) and the number of binding motifs normalised by the length of the genome (right side figure). The gray lines connect the same virus species and serotypes. The abbreviations used for the viruses: Jap (Japanese encephalitis), Nile (West Nile Fever), Chiku (Chikungunya), Yellow (Yellow Fever) and Tick (Tick-borne encephalitis).

The structural content revealed interesting clustering of the viruses. For example, while having a different genome sequence, Dengue viruses also cluster together with Ebola, since they share a less structured genome. Interestingly, DENV-2, the most virulent and spread serotype ^7,8^, is similar in structural content to Ebola, probably the most lethal viral hemorrhagic fever ^28^. According to its structural content, Zika is also part of the sub-cluster, together with West Nile Fever and Japanese Encephalitis. The mosquito-transmitted Yellow Fever and Chikungunya form a cluster, indicating that their structural content is similar. This is partially in agreement with the VHFs tree ^27^. Interestingly, since Tick-borne Encephalitis is more structured than any of these viruses, it forms an outlier. This is not a surprise, since it was previously shown that the secondary structure of mosquito- and tick-borne flaviviruses s are more different, especially in the 3’ UTR ^32^. To conclude, these results indicate that the level of secondary structure inside a viral genome can be used as a metric to build a tree of similarities, which could be further employed to classify viruses.

### Interaction between viral genomes and human host proteins can be used to classify VHFs

During translation and replication, ss-RNA viruses are naked RNA molecules inside human host cells. Previous studies already showed that genomes of the Dengue viruses interact with multiple human proteins during the infection and that the protein binding can enhance or inhibit the virulence ^33^. Furthermore, RNA binding proteins tend to exhibit an altered activity during viral infection, in some cases due to the presence of highly abundant viral RNA, which can compete for the interaction with cellular RNA ^34^. To study the relationship between human proteins and the viral RNA structures, we selected binding motifs from RNA Bind-n-Seq (RBNS) data of 78 human RNA-binding proteins ^23^, and searched the complete viral genomes for these motifs (Methods: Protein-RNA interactions). We observed that the 4 dengue serotypes have a different presence of protein binding domains, with DENV-2 showing the highest number of motifs, followed by DENV-4 (Supplementary Figure 1).

Similarly to the structural content analysis above, we used the number of protein binding domains to classify the viruses. To further understand how the connection between structure and interaction with proteins can classify viruses, we compared the resulting trees (Figure 4). Interestingly, the Dengue cluster is almost perfectly maintained, except that, for the number of protein interactions, DENV-2 is more similar to Chikungunya than Ebola, which in turn is more related to Yellow Fever. Furthermore, clustering of West Nile Fever, Japanese Encephalitis and Zika is partially maintained when using structure and interaction with proteins. To conclude, by analyzing thousands of different viral genomes, we identified specific clusters, conserved both in terms of secondary structure content and the potential number of interactions with proteins.

### Relationship between the structural content and number of protein interaction motifs

Since both the structural content and the number of binding motifs could be used to classify the viruses, we hypothesized that there is a correlation between these two features. We found an overall high anti-correlation (r=-0.74; p-value < 2.2e-16) between the number of protein-binding motifs and the structural content in Dengue, meaning that less structured Dengue viruses tend to bind more proteins (Figure 5a). Interestingly, the different dengue serotypes cluster together according to their structure and the interaction with proteins (Figure 5a). Also, the serotypes show a different trend when independently analysed, with DENV-3 and DENV-4 exhibiting the highest influence of the structure on the number of possible interacting proteins (r is -0.31 and -0.36; p-value < 2.2e-16 and 7.177e-07, respectively; Figure 5b).

**Figure 5.**
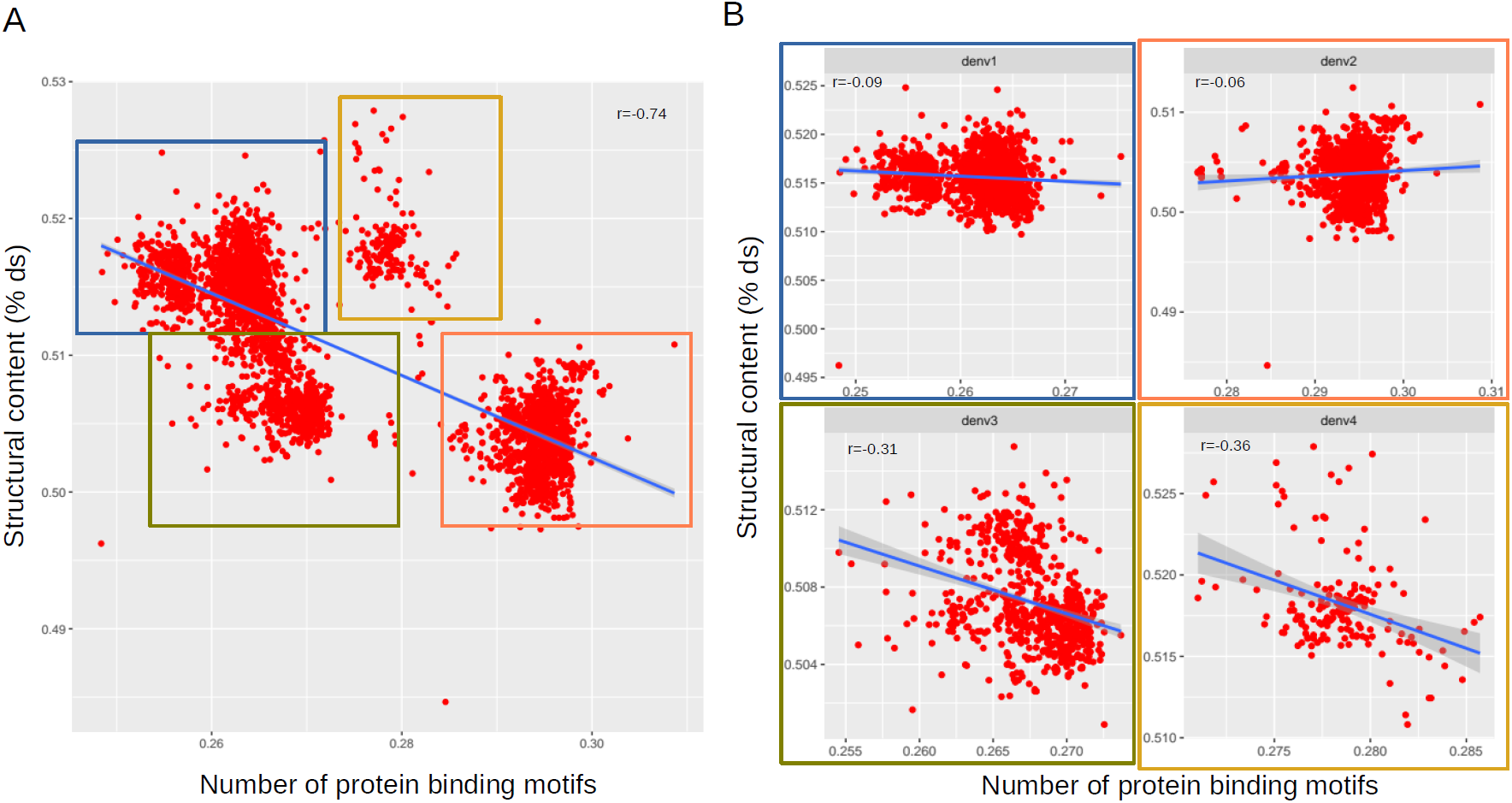
Correlation between the secondary structure and the interaction with proteins for all the dengue genomes. (A) Correlation between the structural content and the averaged number of binding domains for all the dengue genomes. DENV1 is indicated by a blue box, DENV2 by an orange box, DENV3 by a green box, and DENV4 by a golden box. (B) Correlation between the structural content and the averaged number of binding domains independently for the 4 dengue serotypes. The correlation is different when considering each serotype individually.

Next, we compared the secondary structure and protein binding motifs of the other VHFs. The general picture is quite complex, with some viruses showing opposite trends between structure and interaction with proteins, providing a characteristic signature to further classify viruses into three categories. First, similarly to dengue, the mild hemorrhagic fever Chikungunya shows a high anticorrelation (r=-0.84; p-value < 2.2e-16, Figure 6a), identifying a category of less structured but highly interactive viruses. DENV-3 and DENV-4 act similarly to Chikungunya, having less structured genomes but highly interactive with proteins (Figure 5b). Second, Tick-borne Encephalitis and Zika show a positive correlation (Pearson correlation of 0.23 and 0.89; p-value 0.01 and <2.2e-16 respectively; Figure 6), identifying a category of high-structured viruses that are also highly interactive with host-proteins. Third, a class composed of Japanese Encephalitis, West Nile Fever, and Ebola show almost no correlation between the structure and the possible interaction with proteins (r∼0). These results indicate a complex relationship between the structural content and protein-binding motifs, which can classify the viruses in 3 different categories: highly-structured and highly-interactive, poorly-structured and highly-interactive, or without a strong relationship between the overall structural content and interaction with proteins.

**Figure 6.**
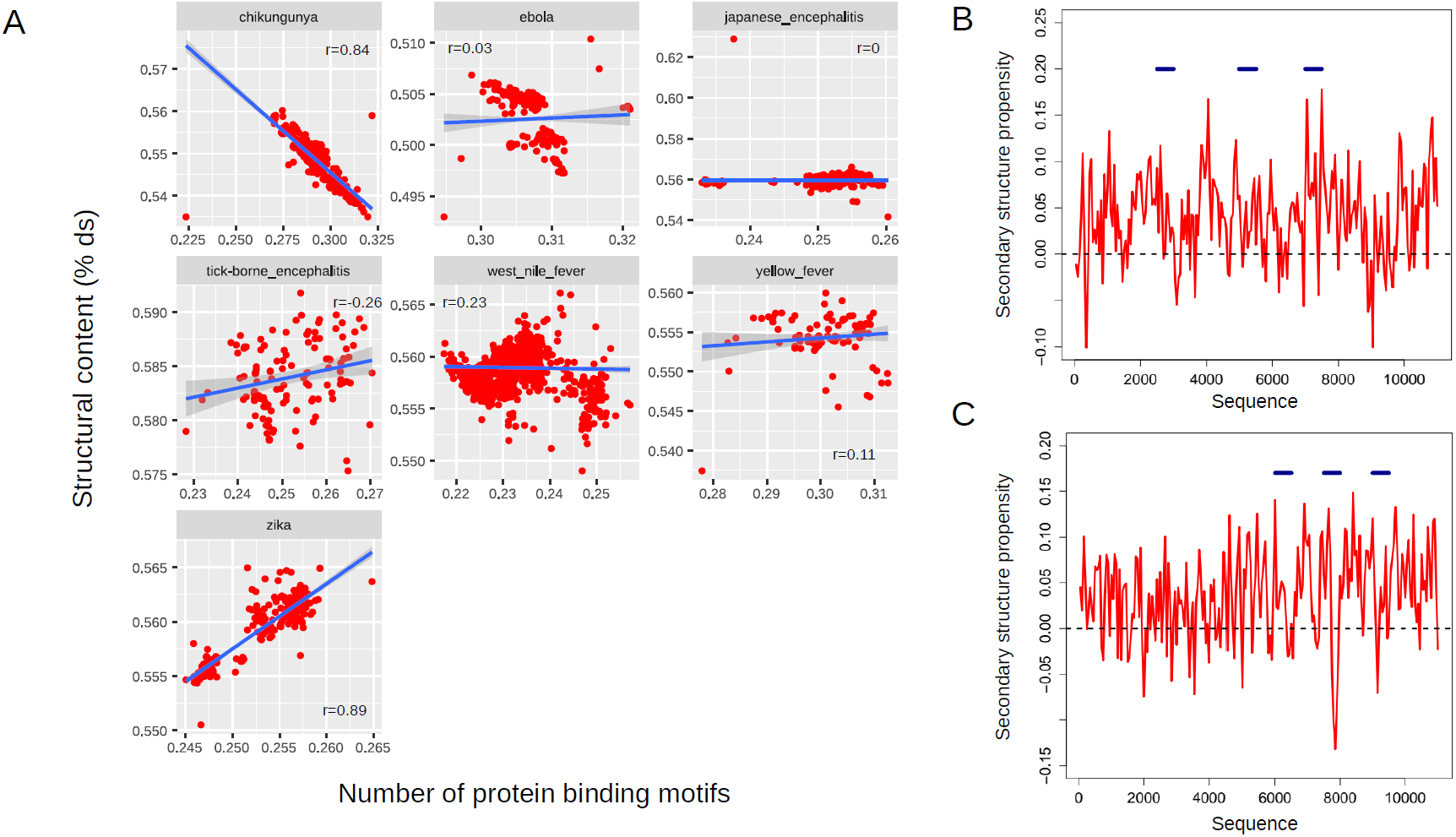
Correlation between the structural content and the averaged number of binding domains for the remaining 7 viral species. (A) Each point represents a single genome, while x- and y-axes indicate its structural content and the number of protein binding domains divided by the averaged size of the genome, respectively. (B) Consensus profile obtained from the secondary structure profiles of all the Zika genomes. Regions considered double-stranded for the majority of Zika genomes have a propensity >0, while a score <0 identifies consensus single-stranded regions. The blue bars mark the top 3 regions of 500 nt with the highest count of protein binding motifs. (C) Consensus profile obtained from the secondary structure profiles of all the Chikungunya genomes. Regions considered double-stranded for the majority of Chikungunya genomes have a propensity >0, while a score <0 identifies consensus single-stranded regions. The blue bars mark the top 3 regions of 500 nt with the highest count of protein binding motifs.

To further understand the different behaviour of Zika and Chikungunya in terms of secondary structure and interaction with proteins, we generated a consensus-profile between the hundreds of genomes available (Methods; Figure 6b, c). We also highlighted for each species the 3 regions in windows of 500 nt which have the highest-presence of protein binding motifs (Figure 6b, c; blue bars). Interestingly, this analysis validate our overall observation, where the most contacted regions in Zika are very structured in the consensus profile obtained from all Zika viruses (Figure 6b). Moreover, the most structured region from Zika consensus profile and one of the most highly-contacted by proteins is the one encoding for the nonstructural protein 3 (NS3), a helicase essential for viral replication ^35^. We speculate that this region needs a very specific structure in order to be highly regulated by proteins. Conversely, the most contacted regions for Chikungunya consensus secondary structure profile fall into highly unstructured regions (Figure 6c). The least structured region of Chikungunya consensus secondary structure profile, and one of the most regulated by proteins, encode for the structural protein E3 ^36^.

To extend and further validate this result, we checked the interactions between the most structured Zika and Chikungunya viruses and >1000 human RBPs. After selecting the 10 most and least structural genomes of Zika and Chikungunya, we used the catRAPID algorithm ^24^ to predict > 4×10^6^ protein interactions between the genomes and human proteins (Methods: Protein-RNA interactions). Interestingly, the highly-structured Zika viruses have stronger and more frequent interactions with proteins, reaching x10 more strong interactions with proteins, compared to Chikungunya (Supplementary Table 1). This result supports our hypothesis that Zika interacts with human proteins mainly using double-stranded regions when compared to Chikungunya.

### Geographical distribution of Dengue serotypes

We also studied the geographical connection between Dengue serotypes and the secondary structure content. We found that the African DENV-1 is predicted to be more structured than the other serotypes, even when compared with the Asian strains (Supplementary Figure 23 Kolmogorov-Smirnov = 0.001), while on the contrary, Asian DENV-3 is more structured than the African strains (Kolmogorov-Smirnov = 0.07; Supplementary Figure 3).

The correlation between binding motifs and secondary structure at the geographical level portrays a quite complex scenario (Figure 7). The clearest clusters are identifiable for DENV-3, where the South-American strains are not only less structured, but also highly interacting with proteins. DENV-3 from South-America is also more distant from the Asian strains (Euclidean distance centroids x1000=5), compared, for example, with DENV-1 (Euclidean distance centroids x1000=1).

**Figure 7.**
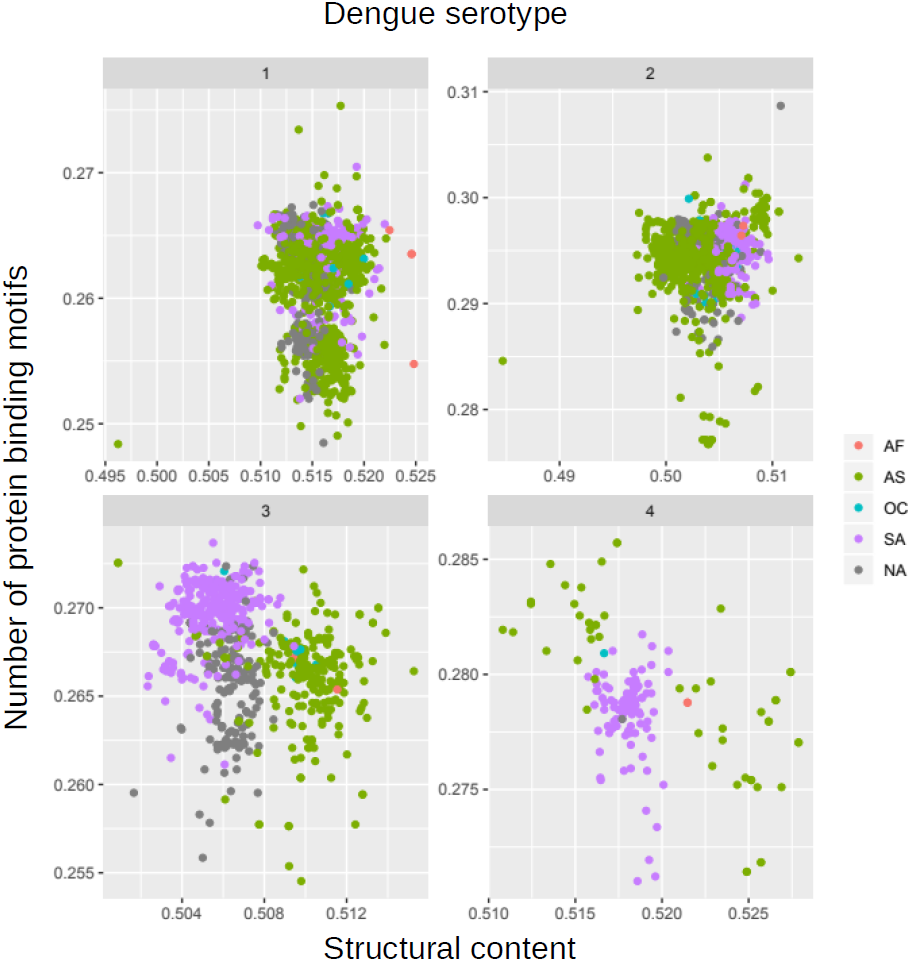
Correlation between the structural content and the averaged number of binding domains for the dengue serotypes, according to their geographical distribution. Samples coming from Africa (AF), Asia (AS), Oceania (OC), South America (SA), and North America (NA) were marked using different colours. Each point represents a single genome, while x- and y-axes indicate its structural content and the number of protein binding domains divided by the averaged size of the genome, respectively.

## DISCUSSION

The genomes of viral hemorrhagic fevers show different level of secondary structure, especially structured in the UTR regions ^14,16^. This secondary structure is thought to be needed for different viral mechanisms, such as packaging and egression ^11–13^. However, a comprehensive secondary structural landscape for their genomes was lacking. In our work, we computationally profiled and analysed the secondary structure profiles of more than 7500 complete viral genomes, including almost 4000 dengue samples, and 3500 other viral hemorrhagic fever-causing viruses. By studying the structural profiles, we observed that Dengue viruses are predicted to be less structured compared to viruses such as Zika, Yellow Fever, and West Nile Fever (Figure 2). Conversely, Dengue serotypes still tend to retain structured UTRs, probably to be efficiently translated by the cellular machinery, similarly to human mRNAs ^26,29^. Single-stranded regions could be necessary to confer flexibility to the viral genomes, since flaviviruses need a high level of structural plasticity to undergo conformational changes during their life cycle, including circularization ^37^.

We also identified a correlation between the secondary structure and the number of protein binding domains, implying that the secondary structure is employed to regulate potential binding with proteins, as observed for human RNAs ^38^. For viruses, the situation is more complex, as we observed a significantly positive (Tick-borne Encephalitis and Zika) and negative (Chikungunya, DENV-3, and DENV-4) relationship between secondary structure and the potential interaction with proteins. We speculate that the opposite behaviors of the viruses are caused by the fundamental differences between viruses. For example, Zika (positive RNA genome, positive correlation between secondary structure and interaction with proteins) and Chikungunya (negative genome, negative correlation) belong to different viral families (Flavivirus and Togavirus respectively), and have a completely different capsid. Also while Zika shows a sequence similarity of 56% with Yellow Fever and Japanese encephalitis, it shows only 1.3% sequence identity with Chikungunya ^39^.

We also analysed the secondary structure at a geographical level, showing that DENV-3 strains from South-America and Asia have different patterns in their structure and the potential interaction with proteins, especially when compared with DENV-1 and DENV-2 (Figure 7). Interestingly, this is in line with DENV-3 being the youngest serotype and the only one with a proposed origin not in Asia but in America ^40^. This could explain the niched behaviour of DENV-3 in terms of structure and protein interactions, as well as supporting a possible independent origin of Asia and American DENV-3. Interestingly, DENV-4 shows a similar trend, but there are too few samples available to explain its evolution.

## CONCLUSIONS

To our knowledge, this is the first study that employed secondary structural content and the presence of protein-binding domains to build similarity trees between VHFs. The secondary structure and interaction with proteins can be used to cluster the viruses in agreement with previous phylogenetic trees, such as Dengue serotypes, and Japanese Encephalitis with West Nile Fever. Conversely, some relationships are surprising, as for example, DENV-2 is closer to Ebola when the secondary structure is used to establish similarity, but not when using the interaction with proteins. This result suggests how different measures, especially the secondary structure content, could be used to classify further and characterise different classes of viruses.

Our massive computational analysis provided novel results regarding the secondary structure and the interaction with human proteins, not only for Dengue serotypes but also for other viral hemorrhagic fevers. We envision that these approaches can be used by the scientific community to classify further and characterise these complex viruses.

## ABBREVIATIONS

CROSS: Computational Recognition of Secondary Structure
SHAPE: Selective 2′ Hydroxyl Acylation analyzed by Primer Extension
PARS: Parallel Analysis of RNA Structure
VHF: Viral Haemorrhagic Fevers
DENV: Dengue virus
RBP: RNA binding protein
UTR: Untranslated region
CDS: Coding sequence
AUC: Area the ROC curve
ss-RNA: single-stranded RNA
MFE: Minimum free energy

## SUPPLEMENTARY FIGURES

**Supplementary Figure 1**. ROC curves of our predictions obtained using CROSS and experimental SHAPE data on DENV-2 (A) and DENV-1 (B). SHAPE data were ranked according to their reactivity, and the 5%, 10% and 25% top/bottom nucleotides were selected. The AUC increases from 0.75 (25% top/bottom ranked SHAPE data; i.e. half of the dataset) to 0.85 (5% top/bottom ranked SHAPE data). Uncharacterised SHAPE reactivities <0 were removed from the ranking.

**Supplementary Figure 2**. Violin plot showing the interaction with proteins for each dengue serotype, computed as the presence of RNA binding motifs on their genome, averaged for the mean of the length of each serotype.

**Supplementary Figure 3**. Barplot showing for each dengue serotype the differences in structural content (% double-stranded nucleotides) in different geographical samples coming from Africa (AF), Asia (AS), Oceania (OC), South America (SA) and North America (NA).

**Supplementary Table 1**. Number of predicted interactions of all the human proteome and the 10 most structured Zika and Chikungunya genomes. An increasing threshold on the Discriminative Power (DP) of catRAPID algorithm was used to iteratively select stronger interactions.

## DECLARATIONS

### Ethics approval and consent to participate

Not applicable.

### Consent for publication

Not applicable.

### Availability of data and materials

Not applicable. All the data we used were available in public datasets.

## Author’s contribution

RDP conceived the study. MM and RDP designed the study. RDP performed the analysis. RDP and MM wrote the manuscript.

## Acknowledgments

The authors thank the other members of Mutwil’s group for useful comments.

