## Supplementary figures and images for "Structural landscape of the complete genomes of Dengue serotypes and other viral hemorrhagic fevers"

### Supplemental Figures

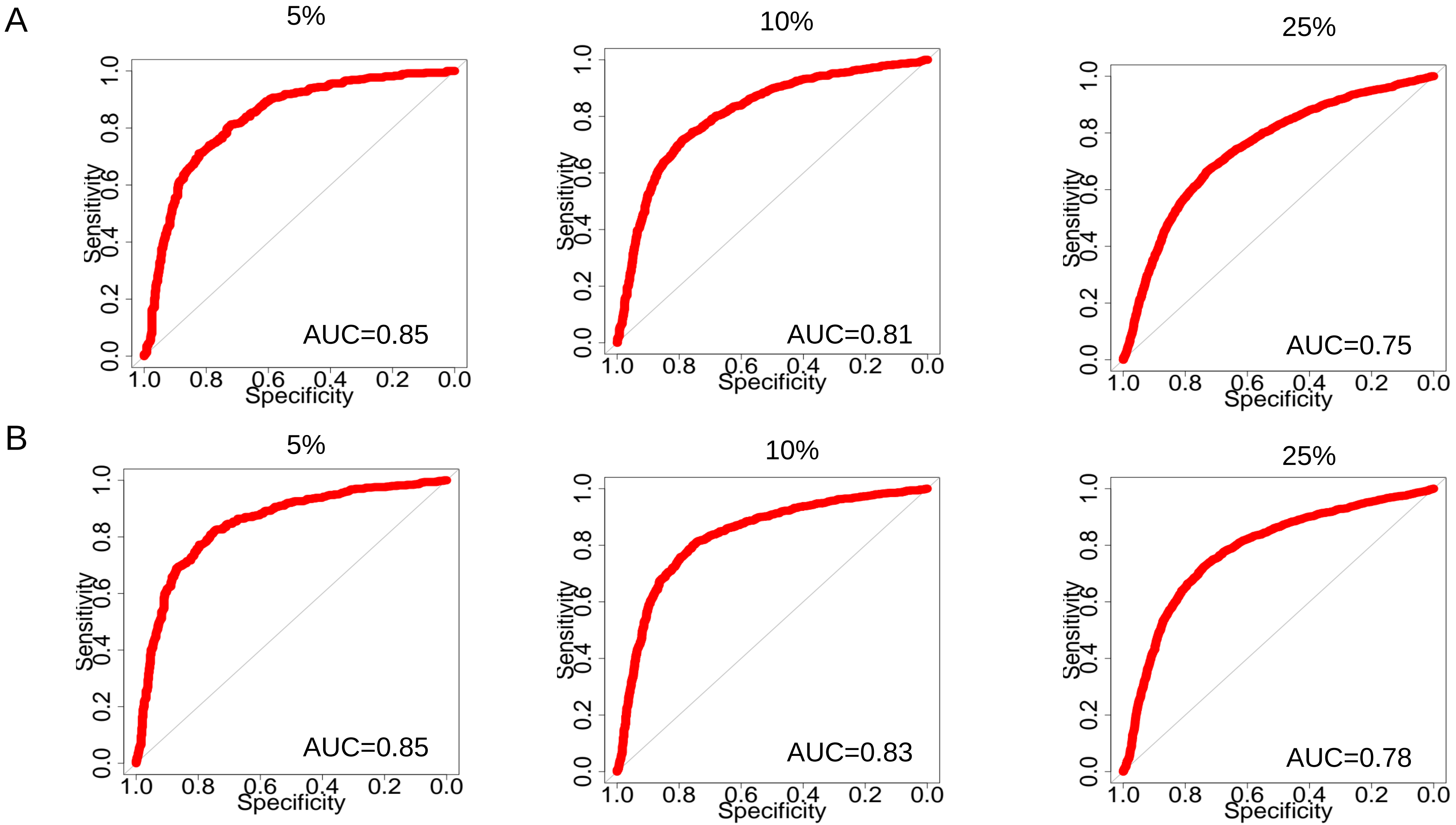

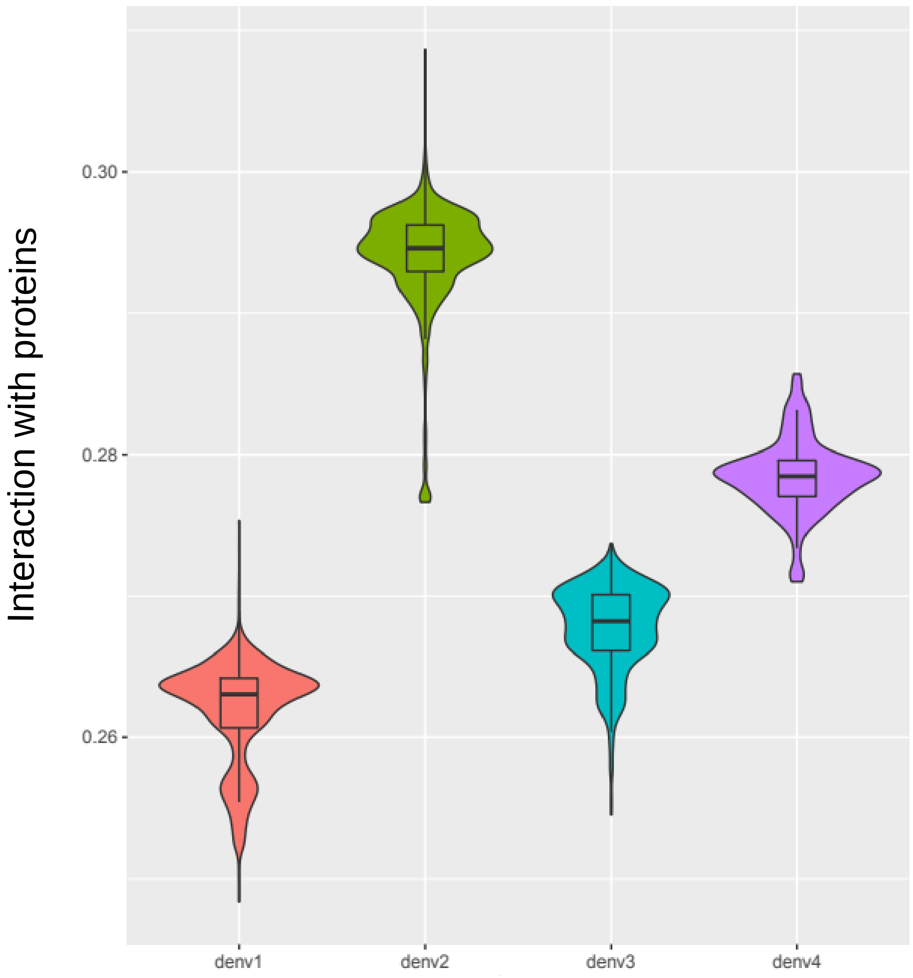

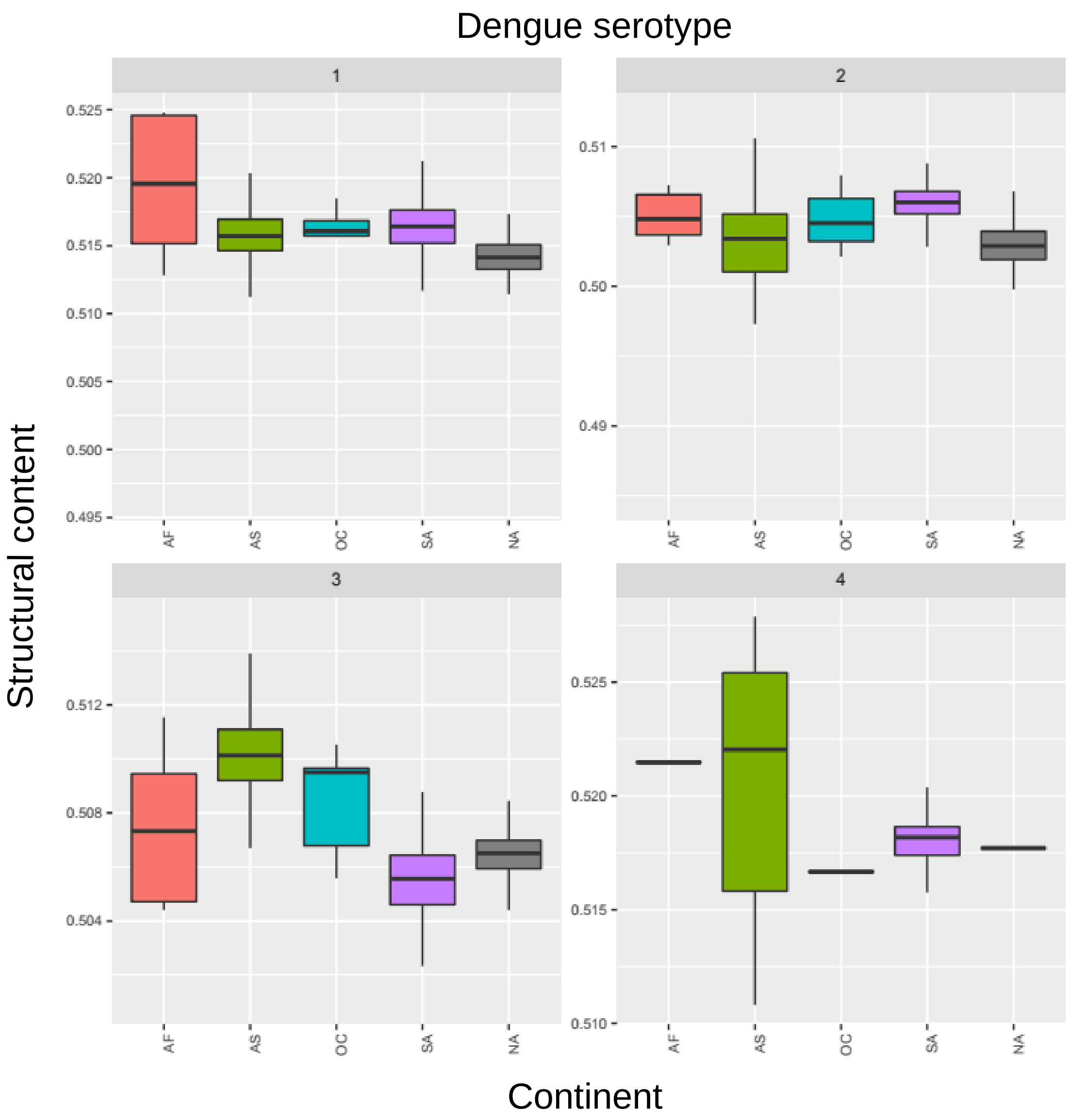

| | DP>0.5 | DP>0.7 | DP>0.9 |
|---|---|---|---|
| **Zika** | 234’816 (x2.5) | 80’017 (x4) | 8’546 (x10) |
| **Chikungunya** | 90’909 | 19’056 | 773 |
